# PathoGFAIR: a collection of FAIR and adaptable (meta)genomics workflows for (foodborne) pathogens detection and tracking

**DOI:** 10.1101/2024.06.26.600753

**Authors:** Engy Nasr, Anna Henger, Björn Grüning, Paul Zierep, Bérénice Batut

## Abstract

**Background:** Food contamination by pathogens poses a global health threat, affecting an estimated 600 million people annually. During a foodborne outbreak investigation, microbiological analysis of food vehicles detects responsible pathogens and traces contamination sources. Metagenomic approaches offer a comprehensive view of the genomic composition of microbial communities, facilitating the detection of potential pathogens in samples. Combined with sequencing techniques like Oxford Nanopore sequencing, such metagenomic approaches become faster and easier to apply. A key limitation of these approaches is the lack of accessible, easy-to-use, and openly available pipelines for pathogen identification and tracking from (meta)genomic data.

**Findings:** PathoGFAIR is a collection of Galaxy-based FAIR workflows employing state-of-the-art tools to detect and track pathogens from metagenomic Nanopore sequencing. Although initially developed to detect pathogens in food datasets, the workflows can be applied to other metagenomic Nanopore pathogenic data. PathoGFAIR incorporates visualisations and reports for comprehensive results. We tested PathoGFAIR on 130 samples containing different pathogens from multiple hosts under various experimental conditions. For all but one sample, workflows have successfully detected expected pathogens at least at the species rank. Further taxonomic ranks are detected for samples with sufficiently high Colony-forming unit (CFU) and low Cycle Threshold (Ct) values.

**Conclusions:** PathoGFAIR detects the pathogens at species and subspecies taxonomic ranks in all but one tested sample, regardless of whether the pathogen is isolated or the sample is incubated before sequencing. Importantly, PathoGFAIR is easy to use and can be straightforwardly adapted and extended for other types of analysis and sequencing techniques, making it usable in various pathogen detection scenarios. PathoGFAIR homepage: https://usegalaxy-eu.github.io/PathoGFAIR/

## Introduction

Foodborne pathogens pose a significant threat to public health worldwide, causing millions of cases of illness and even death every year [1, 2]. These diverse microorganisms, spanning bacteria, viruses, parasites, and fungi, can contaminate a variety of foods, leading to both localised outbreaks and widespread epidemics. Ensuring food safety and controlling foodborne pathogens are key priorities for public health authorities at local, regional, and global levels, including agencies such as the European Food Safety Authority (EFSA), European Centre for Disease Prevention and Control (ECDC), and the World Health Organisation (WHO) [3].

Traditional methods for identifying the source of food contamination require isolation of the target pathogen. This process is not only time-consuming but can be labor-intensive, often requiring multiple steps and sophisticated techniques, and lacks a guaranteed success rate [4]. In contrast, shotgun metagenomic approaches provide a solution to these challenges, as they give an overview of the genomic composition in the sample, including the food source itself, the microbial community, and any possible pathogens and their complete genetic information [5]. Importantly, shotgun metagenomic approaches eliminate the need for prior isolation of the targeted pathogen, as required by Whole Genome Sequencing (WGS) methods, and they are not limited to specific genes as opposed to real-time PCR approaches [6] or 16S rRNA sequencing. While 16S rRNA sequencing is widely used for bacterial taxonomic profiling, it is limited in scope compared to shotgun metagenomic sequencing. The latter allows for the detection of a wide range of pathogens, including bacteria, viruses, and fungi, and gives access to the full genomes enabling the taxa-agnostic identification of antimicrobial resistance (AMR) and virulence genes. This broader scope makes shotgun sequencing more suitable for comprehensive pathogen detection, especially in complex foodborne outbreak investigations [7].

Nanopore sequencing provides long-read data that can capture comprehensive genetic information. Its utilisation, as exemplified by studies like [8], demonstrates its utility in closing genomic gaps, delivering real-time sequencing data, and enhancing the capabilities of metagenomic approaches for outbreak investigations. This technology enables more accurate and rapid pathogen detection, a critical advancement in scenarios where timely responses are essential for effective outbreak management.

Once sequencing data is generated, it must be processed using bioinformatics tools to identify pathogens, their genetic variations, and Virulence Factor (VF) genes, thereby facilitating timely and accurate detection [9, 10]. However, available tools and workflows require bioinformatic and computational knowledge and expertise. For example, tool parameters need to be adapted to the specific use case. End-to-end platforms (Table 1) that allow users to analyse their samples are either restricted with only a limited free trial (e.g. BugSeq [11]) or paid subscription (e.g. OneCodex [12]), or require high computational resources (e.g. SURPI [13] and Sunbeam [14]). For certain free resources, the underlying workflow is not available and adaptable for the user. For example, IDseq [15] (also known as CZID [16]), a free cloud-based service for pathogen detection can only be externally accessed through the dedicated online user interface. Furthermore, some of these workflows are specific to a certain host, pathogen, or sequencing technique, lacking the flexibility for customisation.

**Table 1.** Comparison of features between PathoGFAIR and other similar pipelines or systems. This comparison sheds light on various features and characteristics, such as accessibility, technical specifications, and the scope of analyses offered by each system. It serves as a reference to evaluate the suitability of PathoGFAIR and other similar pipelines or systems for specific needs and requirements

| Features | PathoGFAIR | IDseq | BugSeq | SURPI | OneCodex | Sunbeam | Innuendo<br>[21] | PAIPline<br>[22] | Victors<br>[23] |
| --- | --- | --- | --- | --- | --- | --- | --- | --- | --- |
| **General Characteristics** |  |  |  |  |  |  |  |  |  |
| Free of Charge | ✓ | ✓ | X* | ✓ | X | ✓ | ✓ | ✓ | ✓ |
| Open Source Code | ✓ | ✓ | X | ✓ | X | ✓ | ✓ | ✓ | X |
| Web Interface | ✓ | ✓ | ✓ | X | ✓ | X | ✓ | X | ✓** |
| Automatable API | ✓ | X | X | X | X | ✓ | ✓ | X | X |
| **Accessibility and Availability** |  |  |  |  |  |  |  |  |  |
| Simple end-user Modification | ✓ | X | X | X | X | ✓ | X | ✓ | X |
| Publicly Available Web-server | ✓ | ✓ | ✓ | X | ✓ | X | X | X | ✓ |
| Last Updated | 2024 | 2024 | 2024 | 2014 | 2023 | 2024 | 2018 | 2018 | 2019 |
| **User Support and Documentation** |  |  |  |  |  |  |  |  |  |
| Tutorial | ✓ | X | X | X | X | X | X | X | X |
| Documentation | ✓ | ✓ | ✓ | ✓ | ✓ | ✓ | ✓ | ✓ | ✓ |
| User support | ✓ | ✓ | ✓ | X | ✓ | X | X | X | X |
| **Technical Specifications** |  |  |  |  |  |  |  |  |  |
| Workflow Manager | Galaxy | - | - | - | - | Snakemake | Nextflow | - | - |
| Sequencing Technique | Nanopore*** | Illumina<br>& Nanopore | Illumina<br>& Nanopore | Illumina | - | Illumina | Illumina | Illumina | - |
| **Analyses** |  |  |  |  |  |  |  |  |  |
| Preprocessing | ✓ | ✓ | ✓ | ✓ | ✓ | ✓ | X | ✓ | X |
| Taxonomy Profiling | ✓ | ✓ | ✓ | ✓ | ✓ | ✓ | X | ✓ | X |
| Gene-based Pathogen Identification | ✓ | ✓ | ✓ | ✓ | ✓ | ✓ | ✓ | ✓ | ✓ |
| Allele-based Pathogen Identification | ✓ | X | ✓ | X | X | X | ✓ | X | ✓ |
| Samples aggregation and Visualisations | ✓ | ✓ | X | X | ✓ | ✓ | X | X | X |

\* Free trial of 10 samples is available.
\*\* Malfunctioned when tested.
\*\*\* Can be easily adapted to any other types of sequencing techniques via Galaxy, a customisable and automatable API.

Galaxy [17] is an open-source platform for FAIR data analysis. It enables users to apply a comprehensive suite of bioinformatics tools (that can be combined into workflows) through either its userfriendly web interface or its automatable Application Programming Interface (API) for integrating and customising workflows, enhancing user flexibility. It ensures reproducibility by capturing the necessary information to repeat and understand data analyses. Galaxy offers a collection of high-quality pre-built workflows that can be either used directly or are easily adapted to the user’s needs via the Galaxy workflow editor. Galaxy workflows can be executed on any Galaxy server, even on the private Galaxy server, making it suitable also for data where privacy concerns are important. Furthermore, Galaxy via the major public servers [17] freely provides a large computing infrastructure allowing for the execution of computationally challenging workflows, which is often the case for metagenomic analysis.

Here, we present PathoGFAIR, a collection of Galaxy-based workflows for pathogen identification and tracking its presence among (meta)genomics Oxford Nanopore sequencing data. The workflows are openly available on two workflow registries (Dockstore [18] and WorkflowHub [19]). They can be used directly on three major Galaxy servers (usegalaxy.org, usegalaxy.eu, usegalaxy.org.au) or installed in any other Galaxy server. The workflows are created to work agnostically, detecting all pathogens present in the samples without prior knowledge of the target pathogen. As the workflows are created in Galaxy, they can be adapted, e.g. for other sequencing techniques or with various downstream analyses, such as differential expression analysis, or further statistics and visualisations [17]. Workflows are documented and supported by an extensive tutorial freely available via the Galaxy Training Network (GTN) [20]. Overall, PathoGFAIR offers an easy-to-use computational solution that speeds up the process of sampling, detecting, and tracking pathogens. Links to workflows and tutorials can be found on PathoGFAIR homepage: https://usegalaxy-eu.github.io/PathoGFAIR/

## Implementation

### Overview

PathoGFAIR comprises a collection of 5 workflows, implemented in Galaxy (Figure 1). Each workflow serves a specific function and can be executed independently, enabling users to tailor their analysis according to their requirements.

**Figure 1.**
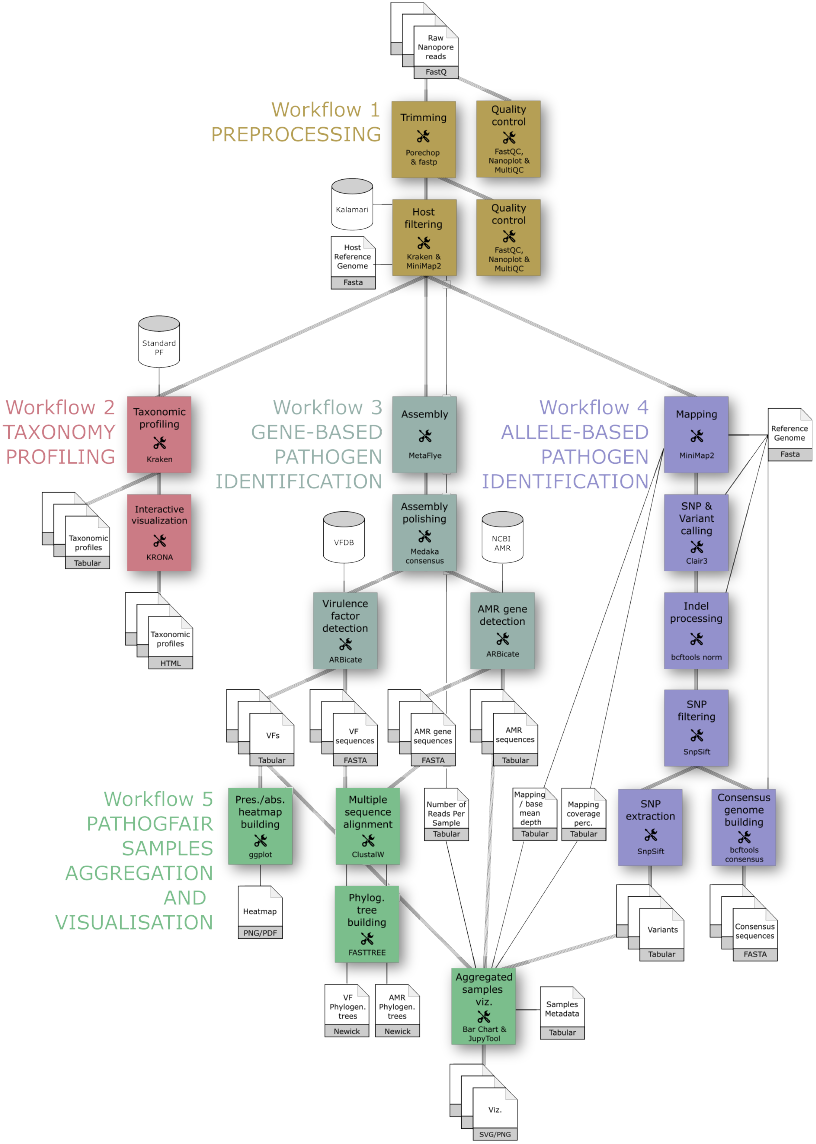
Flowchart of the PathoGFAIR workflows. Workflow 1 (olive green) takes as input sequencing data generated by Oxford Nanopore technologies and performs quality control and host filtering. Then three parallel workflows are executed on the output of Workflow 1: Workflow 2 (red) for taxonomy profiling, Workflow 3 (dark cyan) for gene-based pathogen identification, and Workflow 4 (purple) for SNP-based pathogen identification. These four workflows can run individually and in parallel. Finally, all outputs for the different provided datasets are aggregated in Workflow 5 (green) for PathoGFAIR Samples Aggregation and Visualisation.

The input data for PathoGFAIR comprises sequencing data generated using Oxford Nanopore technologies, along with an optional metadata table describing the datasets. Basecalling for converting raw signal data from the Nanopore sequencer into nucleotide sequences is not included within the PathoGFAIR workflows. In the use cases presented later in the manuscript, real-time basecalling is performed using the MinKNOW software (Oxford Nanopore Technologies) before the reads are used in the workflows. Basecalling is a crucial step, as it affects the quality of the reads. Users are encouraged to ensure that high-quality basecalling is performed before starting the analysis with PathoGFAIR.

The datasets are preprocessed in Workflow 1, which encompasses quality control and host removal procedures. Subsequently, the preprocessed data is directed to three parallel workflows: taxonomy profiling (Workflow 2), gene-based pathogen identification (Workflow 3), and allele-based pathogen identification (Workflow 4). This parallel execution allows for efficient analysis and flexibility in workflow selection. Notably, Workflow 4 can optionally synchronise with Workflow 2 or Workflow 3 to leverage prior taxonomic analysis or gene-based pathogen identification results, providing users with flexibility based on specific use cases. By using detailed taxonomic identification from Workflow 2 or gene-based pathogen identification from Workflow 3, Workflow 4 enhances mapping and SNP detection accuracy. This process involves selecting the correct reference genome of the pathogen for mapping, informed by results from Workflow 2, Workflow 3, or even Workflow 1, which performs initial taxonomy assignment during the host filtering step.

Since each workflow can be executed independently, users can focus on specific aspects of pathogen detection or analysis. This modular approach empowers users to utilise the full range of functions offered by each workflow individually or to combine them as needed for comprehensive pathogen detection.

Finally, in Workflow 5, outputs from the previous workflows and the metadata of the dataset are aggregated and visualised for comprehensive pathogen tracking across samples. This aggregation step ensures a holistic view of pathogen presence and distribution, facilitating further insights and analysis.

Overall, the independent nature of PathoGFAIR’s workflows provides users with a user-friendly and customisable approach to pathogen detection, allowing for both comprehensive analyses and targeted investigations based on specific research needs or objectives.

Ensuring the accuracy and currency of reference data is indeed fundamental for robust metagenomic analysis. PathoGFAIR leverages Galaxy’s integrated Data Managers, which enables Galaxy admins to provide up-to-date reference data. These Data Managers automate the download, installation, and regular update of essential reference databases, ensuring that PathoGFAIR users work with complete, accurate, and up-to-date reference information. PathoGFAIR workflows are configured to use well-maintained and reputable sources, such as NCBI and other public pathogen reference repositories, which further support accuracy and comprehensiveness. Additionally, Galaxy’s user-friendly interface enables users to select preferred references or request the inclusion of specific databases via Galaxy administrators, adding to the workflow’s adaptability for diverse use cases.

PathoGFAIR offers a competitive, and accessible solution (Table 1) to detect and track pathogens in metagenomic Nanopore data through its five Galaxy-based FAIR and customisable workflows.

### Workflow 1: Preprocessing

Workflow 1 encompasses essential preprocessing steps to ensure the quality and integrity of sequencing data.

Quality control and sequence filtering, based on quality, length, or low complexity, are performed using Fastp (v 0.23.2) [24]. Porechop (v 0.2.4) [25] trims low-quality base pairs and removes duplicates and adapters. Quality thresholds are set to ensure that reads have an average quality score of Q20, aligning with accepted standards for Nanopore sequencing, where Q20 or higher quality is typically sufficient for reliable results [26].

Quality-controlled (QC) reads are cleaned of sequences from the host or food source (e.g. bovine in case of bovine meat) by mapping to their reference genome using Minimap2 (v 2.26) (RRID:SCR_018550) [27], a tool tens of times faster than mainstream long-read mappers such as BLASR [28], BWA-MEM [29], NGMLR [30] and GMAP [31] and three times as fast as Bowtie2 [32] designed for Illumina short reads [27]. A variety of reference genomes (e.g. Human, Chicken, or Cow) can be installed on Galaxy servers to work with Minimap2. A wide variety of reference genomes are integrated into Minimap2 on Galaxy, providing users with a convenient selection to choose from before executing the workflow. Kraken2 (v 1.2) [33] is applied for further contamination detection e.g. human sequences using the Kalamari database. The Kalamari database includes mitochondrial sequences of various known hosts [34]. Host/food source reads matched to the Kalamari database are assessed and removed using Krakentools (v 1.2) [33

The workflow returns QC reads without contamination or host sequences as well as interactive reports, produced by FastQC (v 0.12.1) (RRID:SCR_014583), fastp and MultiQC (v 1.11) (RRID:SCR_014982) [35]. Furthermore, Nanoplot (v 1.39.0) [36] is employed to provide detailed quality metrics specifically tailored to the preprocessing step, enriching the suite of analytical insights and facilitating robust data evaluation.

### Workflow 2: Taxonomy Profiling

Workflow 2 performs taxonomic profiling of the microbial community to identify pathogens and other microorganisms for the QC reads from Workflow 1, using Kraken2 (v 1.2) [33] and the PlusPF (archaea, bacteria, viral, plasmid, human, UniVec_Core, protozoa, fungi, and plant) Refseq database (June 7, 2022). Although Kraken2 is a tool designed for short-read sequencing and is known for its false positive taxonomy assignments, particularly at lower microbial abundances [37], its application to longreads can still yield a substantial overview of the microbial community. This is particularly true for discerning bacteria that could potentially be pathogenic at genus and species taxonomic ranks [38, 39]. Kraken2 allows for the rapid assignment of taxonomy at multiple ranks, from kingdom to species, using an efficient exact k-mer matching algorithm. Other tools such as Centrifuge (RRID:SCR_016665) or MetaPhlAn (RRID:SCR_004915) are viable alternatives, also available on Galaxy. Kraken2 is selected for its speed, sensitivity, and ability to work with large reference databases, a critical factor when analysing complex metagenomic samples [33, 40]. The produced community profile is visualised using Krona (RRID:SCR_012785) [41] and observed interactively for different taxonomic ranks using Phinch [42] or Pavian [43].

### Workflow 3: Gene-based Pathogen Identification

In this workflow, the pathogens are identified by the presence of genes associated with pathogenicity. QC reads from Workflow 1 are assembled into contigs using Metaflye (v 2.9.1) (RRID:SCR_017016) [44]. The contigs are then polished using the Medaka Consensus Pipeline (v 1.7.2) [45], which generates consensus sequences using neural networks and shows improved accuracy over graph-based approaches for Oxford Nanopore reads. The polished contigs are afterwards screened using ABRicate (v 1.0.1) [46] for virulence factors (VF) with the Virulence Factor DataBase (VFDB) [47] and for antimicrobial resistance (AMR) genes with AMRFinderPlus [48] database. ABRicate is chosen for its versatility, as it supports multiple databases, including those for antimicrobial resistance genes and the Virulence Factor Database (VFDB). This makes it a comprehensive tool for gene-based pathogen detection, capable of identifying a wide range of relevant genetic markers [46].

### Workflow 4: Allele-based Pathogen Identification

Another approach to identifying pathogens is to use an allelic approach by detecting SNPs, i.e. markers showing evolutionary histories of homogeneous strains [49]. This process includes SNP calling, aimed at identifying novel pathogen strains and elucidating discrepancies compared to reference sequences, thereby facilitating the tracing of emerging variants. Within Workflow 4, both complex variants and SNPs are discerned, serving as crucial elements for subsequent pathogen identification and variant tracing purposes.

QC reads from Workflow 1 are mapped using Minimap2 (v 2.26) to a selected reference genome of a suspected pathogen. Users can choose the reference genome based on their prior knowledge of the target pathogen, the taxonomic analysis in Workflow 2, or the detected pathogenic genes in Workflow 3. Variant calling for mapped reads is performed using Clair3 (v 0.1.12) [50]. Clair3, a tool developed for long reads, has been chosen because it is demonstrated to be faster and more accurate than the Medaka variant pipeline, which its developer has declared deprecated in favour of Clair3 [45]. After that, all complex variants and their information, such as type, genomics position, and quality score, are normalised using bcftools norm (v 1.9) [51]. The normalised reads are filtered using SnpSift filter (v 4.3) (RRID:SCR_015624) [52] based on the SNP quality computed in the SNPs identification step with Clair3. Filtered variants fields required for further analyses are extracted using SnpSift extract fields (v 4.3) (RRID:SCR_015624) [52]. Finally, a consensus sequence for each sample is built using bcftools consensus (v 1.9) (RRID:SCR_005227) [53]. In addition to the variants, this workflow outputs tables including summary metrics like the mapping coverage (breadth of coverage) percentages for every sample, per base covering mean depth (depth of covering), and quality filtered complex variants and SNPs numbers. For more accurate results, users should consider only SNPs with a minimum depth of covering of 10x to ensure reliable calls, as demonstrated in the analyses of the following Use Cases section. This threshold effectively minimises the inclusion of false-positive variants, a challenge often encountered with Nanopore sequencing data due to its inherent error rates.

### Workflow 5: PathoGFAIR Samples Aggregation and Visualisation

In all previously described workflows, individual samples are analysed separately. Workflow 5 consolidates the outputs from Workflows 1, 2, 3, and 4 along with sample metadata to generate various visualisations and reports. These reports illustrate the detected pathogens and facilitate the visualisation and tracking of their presence across all samples.

Virulence Factor (VF) tables from Workflow 3 are used to generate clustered heatmaps showing the VF genes using ggplot2 Heatmap (v 3.4.0) (RRID:SCR_014601). VF sequences are concatenated per sample, generating a consensus sequence of identified VF genes per sample, and aligned over all samples using ClustalW (v 2.1)(RRID:SCR_017277). A phylogenetic tree of the virulence gene sequences is then generated from the multiple sequence alignment using FASTTREE (v 2.1.10) (RRID:SCR_015501) [54] and visualised using Newick Display (v 1.6). The same is performed on the antimicrobial resistance (AMR) tables from Workflow 3. From Workflows 1 and 4 output tables, bar charts are generated.

Other outputs are aggregated and processed within a Jupyter Notebook [55], interactively launched in Galaxy using JupyTool (v 1.0.0). This Notebook showcases the integration of sample metadata to generate analysis-specific plots, leveraging Python (v 3.10.12) [56] libraries such as Pandas (v 1.5.3) [57, 58], Matplotlib (v 3.7.1) [59], Seaborn (v 0.12.2) [60], and Numpy (v 1.24.3) [61]. Examples of these plots include bar plots illustrating the number of reads before and after quality control for all samples, scatter plots visualising relationships between different variables such as pathogen count and sample characteristics, and interactive cluster maps displaying the clustering patterns of samples based on pathogen composition. These visualisation techniques are further elucidated and exemplified in the Use Cases section of this study, where the output tables from the workflows are aggregated with the corresponding sample metadata and visualised to facilitate comprehensive visual analysis.

Virulence factors (VF) and antimicrobial resistance (AMR) genes are often found on mobile genetic elements (MGEs) such as plasmids or phages, meaning they can sometimes appear independently of their bacterial hosts. To address this challenge, PathoGFAIR integrates taxonomic profiling from Workflow 2 with gene detection results from Workflow 3. This cross-referencing ensures accurate attribution of VF and AMR genes to their respective host organisms. For further validation, Workflow 4 enables users to map consensus genomes, generated in Workflow 5 from detected VF genes, against any reference genomes. This process confirms whether these VF or AMR genes, detected in Workflow 3, are genuinely linked to the bacterial genome or merely associated with MGEs, with additional coverage metrics helping to ensure accurate mapping. Future updates to PathoGFAIR will include expanded methodologies to validate gene-host associations using broader taxonomic markers, further refining the precision of pathogen characterisation..

### Workflow Reports

As all PathoGFAIR workflows are designed to run seamlessly on the Galaxy platform, an interactive report is automatically generated upon completion of each workflow. These reports provide a comprehensive overview of the respective workflow’s inputs and outputs. In PathoGFAIR, special attention has been given to refining these reports for enhanced user experience. The reports are carefully curated to automatically showcase and emphasise only the most informative, easily interpretable, and accessible outputs for each workflow. This ensures that users can efficiently extract key insights from the results, facilitating a streamlined and user-friendly analysis experience.

### Easily Adaptable Workflows

The workflows can process raw shotgun (meta)genomics sequencing data from any sample, not only food.

PathoGFAIR has been initially developed to take Oxford Nanopore data as inputs. However, PathoGFAIR can work with Illumina data or other types of sequencing technique data. To adapt to Illumina sequencing only one tool needs to be changed in Workflow 1: Porechop [25] with Cutadapt (RRID:SCR_011841) [62]. Workflows 2, 3, 4, and 5 can be used directly with Illumina datasets without any adaptation. Some tools can be changed based on the tool’s known performance towards short and long reads, such as Clair3 (v 0.1.12) [50] and Metaflye (v 2.9.1) [44]. All the mentioned tools are accessible within Galaxy, allowing for seamless interchangeability.

The workflows can also be adapted to process paired-end reads, by adjusting the tools’ parameters to take paired-end read samples instead of single-end reads. These changes can be applied with little effort by using the user-friendly workflow editor in Galaxy.

Users can seamlessly switch between different host reference genomes and Kraken2 databases, as PathoGFAIR supports various pre-installed databases on the Galaxy servers. This feature enhances user convenience and efficiently explores different configurations to suit specific analysis requirements.

Similarly, tool versions and parameters can be adapted, e.g. to compare results with legacy versions of the workflows. New tool versions are automatically installed on public Galaxy servers using a sophisticated update infrastructure, ensuring a straightforward mechanism to keep the infrastructure up-to-date [63]. Every time a tool is updated, an update of the workflows is suggested, tested with functional tests, and released on the workflow registries once accepted.

Each of the five PathoGFAIR workflows is designed for a distinct type of analysis. Workflows 2, 3, and 4 operate independently, offering the flexibility to run them concurrently or skip them as per user requirements. This modular structure allows users to tailor the analysis to their specific needs, activating only the functionalities necessary for the desired workflow outcome.

### FAIR Workflows

The FAIR principles [64], which emphasise the importance of making research objects Findable, Accessible, Interoperable, and Reusable, offer valuable guidance for optimising the utility and promoting the reproducibility and reusability of any research object (data, software [65], or workflows).

PathoGFAIR has been developed with the FAIR principle in mind and follows the ten tips for building FAIR workflows, as suggested byde Visser *etal*. [66]. First, byusing Galaxyas a workflow manager, the workflows are portable (Tip 6) and come with a reproducible computational environment (Tip 7). The tools integrated into the workflows use file format standards such as FASTA and FASTQ for sequence data, SAM and BAM from the Samtools project for alignment data, VCF for genetic variations, GenBank and GFF3 for genomic annotations, and PDB for structural data (Tip 5) [64]. As explained in the previous section, the workflows are provided with default values (Tip 8) and are modular (Tip 9).

The workflows are available on the GitHub repository of IWC, the Intergalactic Workflow Commission of the Galaxy community (Tip 3) [67]. Workflows in this repository are reviewed and tested using test data before publication and with every new Galaxy release. The IWC automatically updates the workflows whenever a new version of any tool used in these workflows is released. Deposited workflows follow best practices, are versioned using GitHub releases, and contain important metadata (e.g. License, Author, Institutes) (Tip 2). The workflows are automatically added to two workflow repositories (Dockstore [18] and WorkflowHub [19]) to facilitate the discovery and re-use of workflows in an accessible and interoperable way (Tip 1). Via Dockstore or WorkflowHub, the PathoGFAIR workflows can be installed on any up-to-date Galaxy server. They are already publicly available on three main Galaxy servers (usegalaxy.org, usegalaxy.eu, usegalaxy.org.au), which any user can use and modify without restriction.

A thorough explanation of how to use the workflows in PathoGFAIR including a more global description of pathogen identification from Oxford Nanopore data can be found in a dedicated extensive tutorial [68] together with example input data and results (Tips 4 and 10), freely available and hosted via the Galaxy Training Network (GTN) [20] infrastructure.

Finally, for every invocation of the workflows, a Research Object Crate (RO-Crate [69, 70]) can be created to store the data products of the different steps, along with the run-associated metadata (including parameters, tool, and workflow version).

### Use Cases

To showcase PathoGFAIR and its capabilities, 130 samples from two distinct studies—one involving samples with prior pathogen isolation and the other without— were analysed. In the case of non-isolated samples, pathogens were deliberately spiked to mimic real-world scenarios. For isolated samples, prior identification ensured the pathogens’ identities were known. All samples underwent sequencing using Oxford Nanopore technology, highlighting the workflow’s adaptability across diverse sample preparation methods.All workflows of PathoGFAIR were evaluated for their main intended tasks, e.g., the preprocessing workflow for its reads quality retaining and hosts sequences removal performance, but also for their ability to identify the correct pathogen, and how well the accuracy with respect to different sampling conditions is.

### Samples Without Prior Pathogen Isolation

#### Data Generation

In this study, 46 samples had been prepared given the following protocol [71]. Chicken meat was spiked with either one of three *Salmonella enterica* subspecies (*S. enterica* subsp. *houtenae* DSM 9221, *S. enterica* subsp. *enterica* DSM 554, or *S. enterica* subsp. *salamae* DSM 9220) or a mix of them, with concentrations that give Cycle Threshold (Ct) values between 25 and 33. A total of 15 samples were incubated at 37°C for 24 hours before DNA isolation to facilitate bacterial growth. All samples were after incubated at 56°C for 1 hour with lysis buffer and 20 *ng*/µ*l* Proteinase K, followed by DNA extraction according to the STAR BEADS Pathogen DNA/RNA Extraction kit (CYANAGEN SRL, Bologna, Italy) instructions. In this study, approximately 25 mg of meat were used per aliquot for DNA extraction. DNA concentrations were measured with the Qubit® 4.0 Fluorometer (Thermo Fisher Scientific) using the double-stranded DNA (ds-DNA) High-Sensitivity (HS) assay kit (Thermo Fisher Scientific), following the manufacturer’s protocol. The quality was evaluated with a Nanodrop® 1000 (Thermo Fisher Scientific), assessing the 260/280 nm and 260/230 nm ratios. 260/280 and 260/230 ratios were close to the expected ranges 1.8–2.0 and 2.0–2.2, respectively. Extracted DNA was barcoded before sequencing using the Native barcoding genomic DNA (with EXP-NBD104, EXP-NBD114, and SQK-LSK109) protocol (Oxford Nanopore). DNA was then loaded on an R9.4.1 MinION Mk flow cell (Oxford Nanopore). SpotON sample port cover and priming port were closed and sequencing was started. The sequencing device control, data acquisition, and real-time basecalling were carried out by the MinKNOW software of the MinION Mk1C device. For 6 samples, adaptive sampling, a technique used in Nanopore sequencing to selectively sequence microbial DNA while excluding unwanted host DNA (here chicken DNA), was used. Generated sequencing data is available via BioProject PRJNA982679. Metadata for the 46 samples is summarised in Supplementary Table T1 into five pieces of information: (i) expected subspecie(s), (ii) incubation before DNA isolation, (iii) adaptive sampling during sequencing, (iv) Colony-forming unit (CFU)/mL [72], a measure providing a quantitative assessment of viable microbial entities within a given sample and measured using standard microbiological techniques such as serial dilution and plating on agar medium, provides, (v) Cycle Threshold (Ct) values [73], values inversely proportional to the amount of nucleic acid present in the samples.

#### Preprocessing

The number of reads after quality control varies significantly between samples (Figure 2 A), which impacts downstream analyses.

**Figure 2.**
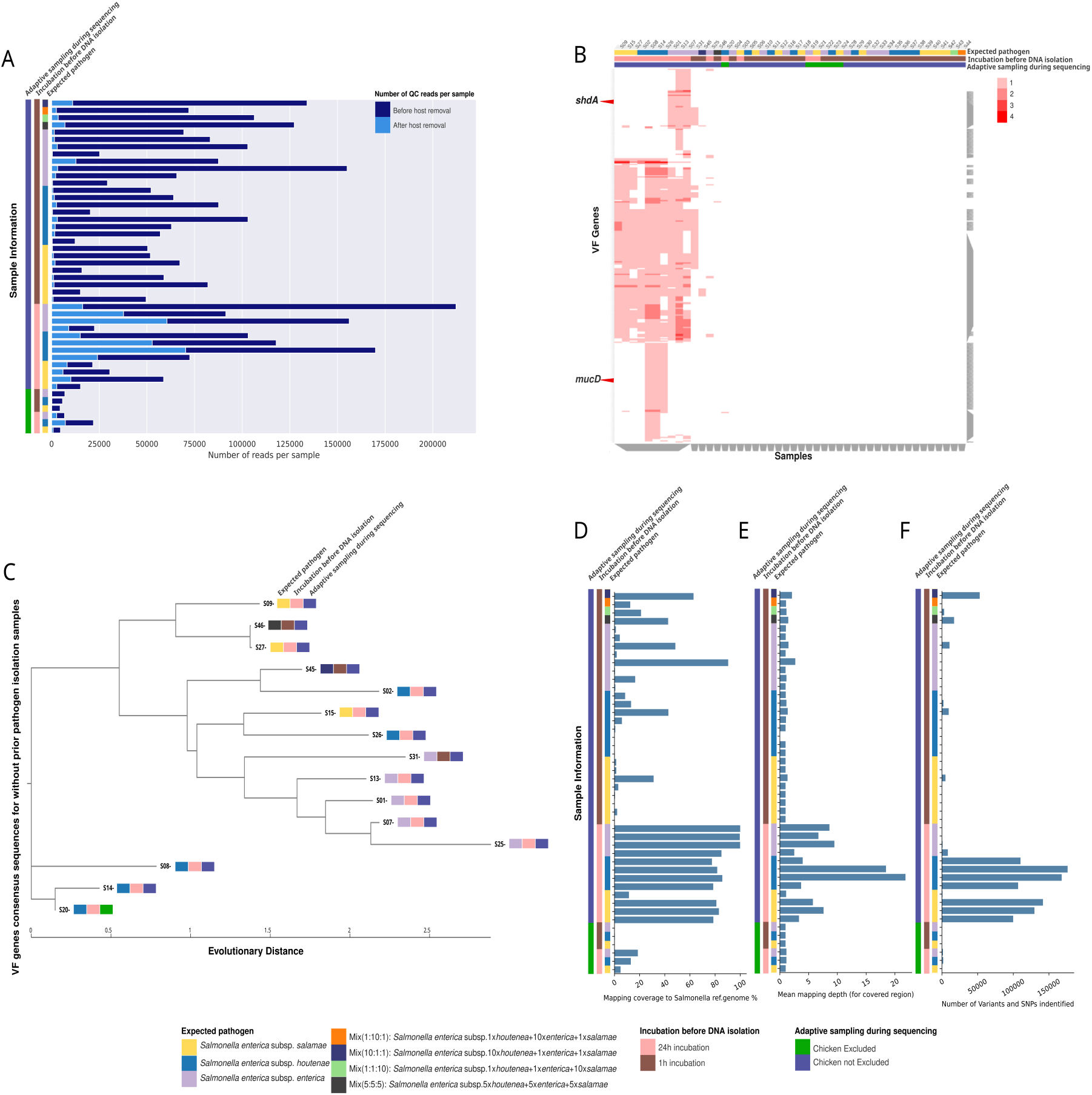
(**A**) Bar plot showing the total number of quality-controlled reads per sample before (dark blue) and after (light blue) host sequences removal. On the left, the metadata of the samples are displayed: (i) the expected *S. enterica* subsp. *salamae* in yellow, *S. enterica* subsp. *houtenae* in blue, and *S. enterica* subsp. *enterica* in light purple), (ii) incubation before DNA isolation (incubated for 24h in pink and incubated for 1h in brown), and (iii) adaptive sampling during sequencing (chicken excluded in green and chicken not excluded in purple)(**B**) Clustergram displaying the identified VF genes’ abundances per sample. The VF genes are presented on the y-axis and all 46 non-isolated samples are on the x-axis along with their sample information. On the top are the metadata of the samples with the same color code as in **A** The grey bars on the bottom and on the right represent dendrogram VF gene (right) and samples’ metadata (bottom) clusters found with hierarchical clustering with a clustering granularity of 0.5. (**C**) Phylogenetic tree, using the nucleotide evolution model; General Time Reversible (GTR) model with a CAT approximation for rate heterogeneity across sites [54]. The Phylogenetic tree was built on the VF genes consensus sequences concatenated per sample and aligned for all samples. (**D**) Bar plot with the mapping coverage (breadth of coverage), i.e. the percentage of covered bases of each sample to the reference genome, measured by calculating the percentage of positions within each bin with at least one base aligned against it. (**E**) Bar plot with the mean of the mapping depth (depth of coverage) of bases mapped to corresponding bases in the reference genome for every sample. (**F**) Bar plot with the number of variants and SNPs found per sample. Mapping coverage percentage and the depth mean indicate whether to trust the variants and SNPs found by the workflow or not, the higher the coverage percentage and the depth mean, the more trusted the SNPs results for the sample.

For host detection using Minimap2 (v 2.26), the option *PacBio/Oxford Nanopore read to reference mapping* was set here. As expected from the samples sequencing protocol (chicken samples and not isolated pathogen), most sequences were assigned to chicken (Gallus gallus galGal6): above 90% in 31 samples and between 55% to 85% for the remaining 15 samples (Supplementary Figure S1). However, the percentage of identified host DNA (between 60% and 98%) was not as low as expected for the 6 samples that had undergone adaptive sampling to exclude chicken DNA during sequencing. This shows that the adaptive sampling to exclude chicken in some samples during sequencing may not have removed all the chicken sequences. All sequences identified as chicken were removed (Figure 2 A). After QC and host removal, 19 samples had less than 1,000 reads. These samples could only be analysed using the taxonomy profiling as highlighted in the next sections.

#### Taxonomy Profiling

*S. enterica* was detected in Workflow 2 for all samples except one, at its species and different subspeciestaxonomic ranks (Interactive KRONA plot in Supplementary online Figure S2 and Supplementary Table T3).

#### Gene-based Pathogen Identification

In Workflow 3, Metaflye (v 2.9.1) tool mode’s option was chosen to be *Nanopore-HQ*, users can expand the workflow and change this option according to their datasets sequencing technique.

No contig was built for 10 of the 27 samples with less than 2,700 reads. The identification of Virulence Factors (VF) or AntiMicrobial Resistance (AMR) genes was then made impossible. For the other 17 samples, only 1 or 2 contigs were created, not enough for identifying VF and AMR genes.

For the remaining 19 samples with created contigs (from 3 to 157) and number of reads higher than 2,700, VF genes were identified in 15 samples (Figure 2 B), 12 of which were incubated before DNA isolation for 24 hours. 3 of the 15 samples were incubated for only 1 hour before DNA isolation, resulting in a few VF genes (Figure 2 B) identified, compared to the other 12 samples, mostly because of the low number of reads (Figure 3 E) from almost the absence of incubation (Figure 2 A). It was, for example, the case for the mixed samples, i.e. samples spiked with all 3 *S. enterica* subspecies, or samples spiked only with *S. enterica* subsp. *houtenae* and adaptively sampled during sequencing.

**Figure 3.**
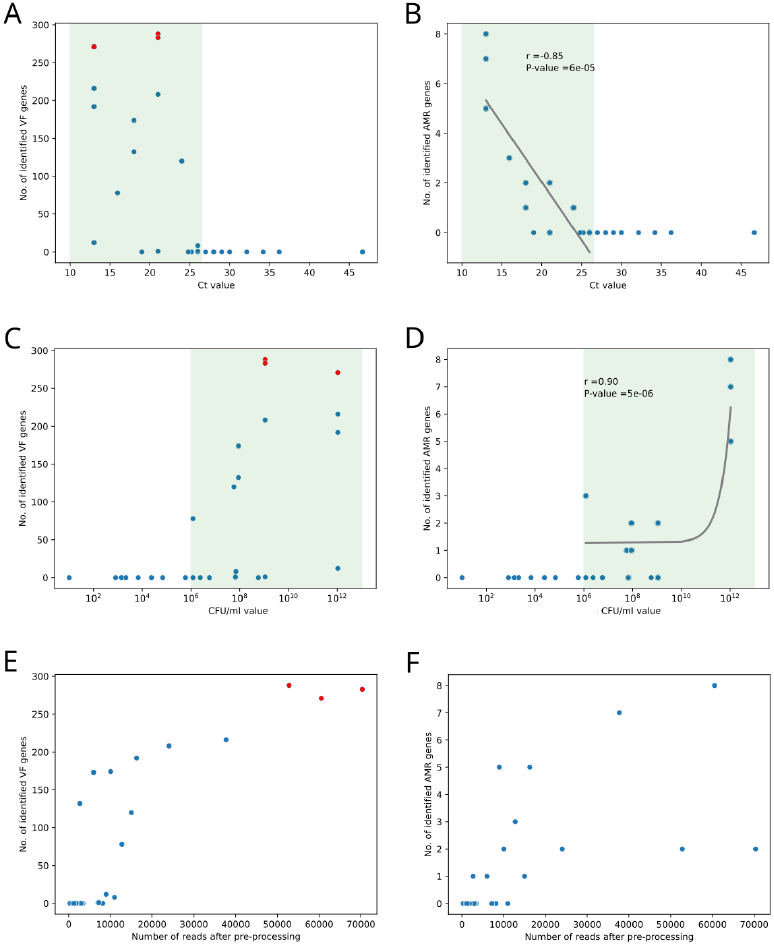
Scatterplots showing the number of identified VF genes (**A, C, E**) and AMR genes (**B, D, F**) in relationship to the Ct value (**A, B**), CFU/mL value (**C, D**), and the number of reads after preprocessing (**E, F**). The green area (**A, B, C, D**) highlights Ct values or CFU/mL values for which genes had been detected. Pearson correlation for values in the green area: (**A**) *R* = –0.33 and *p*-value = 0.23, (**B**) *R* = –0.85 and *p*-value = 5.83*e*^−05^, (**C**) *R* = 0.17 and *p*-value = 0.53, (**D**) *R* = 0.90 and *p*-value = 4.68*e*^−06^

**Figure 4.**
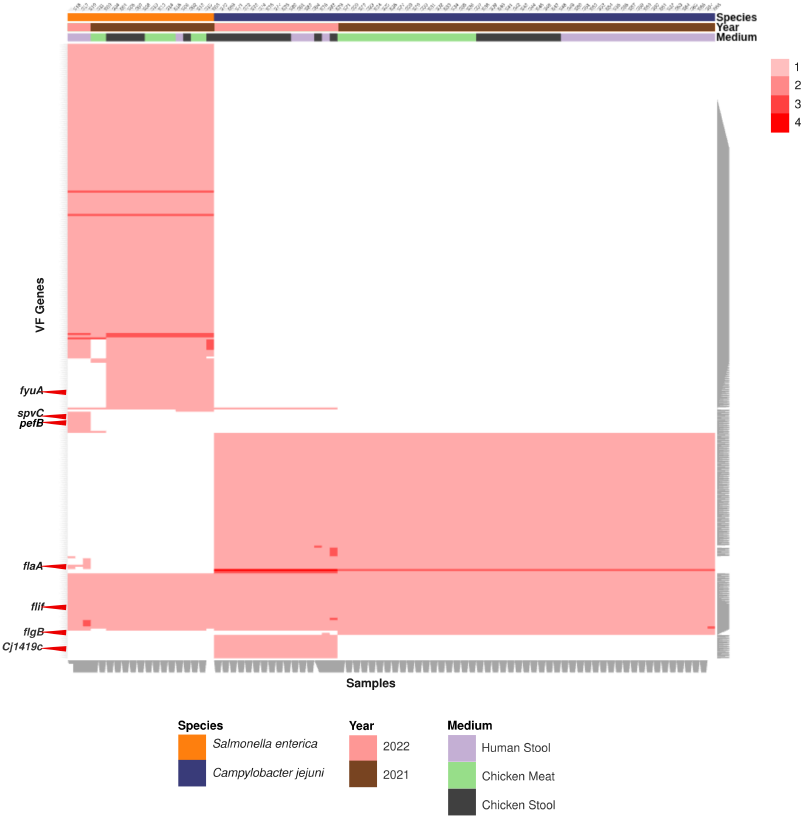
Cluster-map showing the identified VF genes on the y-axis for tested samples presented on the x-axis, clustered based on sample information such as sampling year, isolated pathogen species, and the original host of the sample. Clustering was performed using hierarchical clustering implemented in the Clustergrammer Python package

Some identified VF genes were found more than once in the same sample, with a maximum of 4 times. Common VF genes were identified for samples expecting identical *S. enterica* subspecies (Figure 2 B), such as the *mucD* gene, a serine protease *mucD* precursor, which was only found in *S. enterica* subsp. *houtenae* spiked samples, or *shdA*, an AIDA autotransporter-like protein, only found in *S. enterica* subsp. *enterica* spiked samples, but not in samples spiked with only *S. enterica* subsp. *houtenae* or *S. enterica* subsp. *salamae*.

Similar results were found for AMR genes (Supplementary Figure S3, 3 F). The sampling conditions affected the number of identified VF and AMR genes as shown by the relationships between the Ct value, CFU/mL value, or the number of remaining reads after preprocessing (Figure 3). The lower the Ct value, the higher the number of VF genes and AMR genes identified (Figure 3 A & B). No VF or AMR genes were detected for samples with Ct values above 26. For Ct values below 26, there was a negative correlation (Pearson *R* = –0.85, *p*-value = 6 *×* 10^−05^) between the Ct value and the number of identified AMR genes. Similar but inverse relations were observed for CFU/mL value (Figure 3 C & D), with a threshold for VF and AMR gene detection at 10^6^. VF and AMR genes were then detected if several conditions were fulfilled: a Ct value below 26, CFU/mL value above 10^6^, and at least 5,000 reads after preprocessing. The further the samples were from these thresholds, the higher the number of VF genes and AMR genes identified. Indeed, the three top scattered dots (in red - Figure 3 A, C & E), with identified VF genes between 250 and 300 were the samples with the highest number of reads, higher CFU/mL value, and a relatively lower Ct value compared to other samples. Generally, allowing samples to incubate for a short period before sequencing enhances microbial growth, resulting in higher CFU/mL values and lower Ct values. This increase in microbial concentration improves the efficiency of direct sequencing by providing more genetic material for analysis, facilitating faster and more accurate pathogen detection.

#### Allele-based Pathogen Identification

In Workflow 4, samples were mapped against a reference genome of an expected pathogen chosen by the user. *S. enterica subsp. enterica* ser. Typhimurium (NC_003197.2) was chosen for this data, as it is widely recognised and extensively used in genomic studies due to its complete and well-annotated genome sequence [74]. However, given the diversity among the serovariants of *S. enterica subsp. enterica*, a high number of complex variants and SNPs are anticipated.

The provided mapping statistics (mapping coverage (breadth of coverage) and mapping depth (depth of coverage) in Figure 2 D, E) serve as proxies for assessing the number and quality of identified SNPs (Figure 2 F). SNPs with low mapping depth are less reliable than those with higher depth. Reliable SNP calling typically requires a depth of at least 10, achieved in 2 samples. Samples with the highest mean mapping depth corresponded to samples with the highest number of reads after preprocessing (Figure 2 A). The higher the coverage and the mean mapping depth, the more quality SNPs were identified (Figure 2 D-F). Some of the samples spiked with *S. enterica* subsp. *enterica* had a high breadth of coverage but a low mean depth of coverage depth, as a result, the number of their quality filtered identified SNPs was low.

#### PathoGFAIR Samples Aggregation and Visualisation

For the samples for which VF or AMR genes had been identified, phylogenetic trees were built on the concatenated genes consensus sequences (Figure 2 C for VF genes, Supplementary Figure S4 for AMR genes). These trees help track divergence between samples and could then highlight the contamination point or an evolution of the subspecies because of mutations. Indeed, samples spiked with *S. enterica* subsp. *enterica* were found together in the VFs-based tree (Figure 2 C), so the identified VF genes were unique to these samples and could clearly separate the samples from samples with other *S. enterica* subspecies. The samples spiked with *S. enterica* subsp. *houtenae* were mostly clustered together, except for 2 samples because of extra identified VF genes common with samples spiked with *S. enterica* subsp. *enterica* and/or *S. enterica* subsp. *salamae*. The 2 samples spiked with a mix of the 3 subspecies were found in the middle of the tree (Figure 2 C), showing that a mix of VF genes related to the different subspecies was identified. The mixed sample, S45, spiked with a higher concentration of *S. enterica* subsp. *houtenae* than the other subspecies, was close to the sample, S02, spiked with *S. enterica* subsp. *houtenae* only. For AMR genes phylogenetic tree (Supplementary Figure S4), samples were not as clearly separated as the tree for VF genes, mostly because the number of identified AMR genes was relatively low compared to the number of identified VF genes.

#### Sensitivity

The performance of the workflows was evaluated based on their ability to identify the expected *S. enterica* pathogen, as well as *S. enterica* subspecies and strain taxonomic ranks for the tested samples (Supplementary Table T3). In a metagenomic setting, other detected species cannot be regarded as false positives, as they may naturally be present in the sample. Therefore, only sensitivity was reported.

For the taxonomy profiling (Workflow 2), the expected pathogen was detected at its species taxonomic rank in all but one sample, resulting in a sensitivity of 97.8%. At the subspecies taxonomic rank, the expected subspecies was detected in 28 out of 46 samples, yielding a sensitivity of 64.0%. To further evaluate subspecies classification performance, the sample-wise sensitivity (the percentage of correctly identified *S. enterica* subspecies out of all detected *S. enterica* subspecies) was calculated. Averaged across all samples, the sample-wise sensitivity was 47.3%. In the gene-based pathogen identification (Workflow 3), at least one virulence factor (VF) gene of the expected pathogen, at strain taxonomic rank, was detected in 13 out of 46 samples, corresponding to a sensitivity of 28.2%. For the samples in which no VF gene was detected, no contigs could be generated, preventing gene calling.

Changing the workflow’s default settings, such as using different reference databases for preprocessing in Workflow 1, taxonomy profiling in Workflow 2, or gene-based pathogen identification in Workflow 3, would likely impact these metrics. Different reference databases could influence the accuracy and sensitivity of taxonomic classification and pathogen identification, as they may contain varying levels of strain-specific data. Adjusting parameters like threshold values, filtering criteria, or the inclusion of additional databases could also affect the detection sensitivity and overall performance, potentially improving or reducing the workflow’s ability to accurately identify pathogens and associated genes in the given samples.

### Samples With Prior Pathogen Isolation

#### Data Description

To further test PathoGFair, 84 public datasets were used [75]. These samples were sampled in Palestine by the Swiss Tropical and Public Health Institute either from chicken meat, chicken stool, or human stool, in 2021 or 2022 (Supplementary Figure S5). In these samples, *S. enterica* had been isolated in 19 samples, and *Campylobacter jejuni* in 65 samples. The generated sequencing data are provided under BioProjects PRJNA942086 (S. enterica [76]) and PRJNA942088 (C. jejuni [77]).

#### Preprocessing

Negligible contamination or host sequences were found between 0% and 0.02% (Supplementary online Figure S6), as expected because of the prior isolation of the pathogen. The number of reads ranges between 3k and 217k reads per sample, after quality control.

#### Taxonomy Profiling

As presented in the interactive KRONA plot (Supplementary online Figure S6), the first 19 samples, *S. enterica* isolates, were assigned correctly to *S. enterica*, and the remaining 65 samples were assigned correctly to *C. jejuni*. With the KRONA plot (Supplementary online Figure S6), the total number of reads for each sample can be seen along with detailed percentages on the assigned taxa at each taxonomic rank.

#### Gene-based Pathogen Identification

In this workflow, we identified VF and AMR genes for all samples, thanks to the higher number of reads retained after preprocessing and the prior isolation of the pathogens. Consequently, VF genes were detected in all samples, with more VF genes identified than AMR genes (Supplementary Figure S7), as in general, the number of VF genes in a bacterial genome is often higher than the number of AMR genes. Samples containing *S. enterica* exhibited more VF genes (172 to 207) compared to samples with *C. jejuni* (96 to 113). The opposite trend was observed for AMR genes, *C. jejuni* samples typically had 12 AMR genes detected, while *S. enterica* samples mostly had 6 AMR genes (Supplementary Figure S7)

The analysis revealed that samples with similarly isolated pathogens clustered together based on detected VF genes (4). For example, samples with *S. enterica* and *C. jejuni* formed distinct clusters. Moreover, correlations were observed among samples from different hosts, sampling years, and pathogenic species.

Specific VF genes were found in samples with similar isolated pathogens, indicating potential subspecies-specific differences. For instance, Cj1419c, a methyltransferase Capsule biosynthesis and transport gene product, was exclusively found in *C. jejuni* isolates samples sequenced in 2022, while flgB gene, encoding flagellar basal body rod protein, was only detected in *C. jejuni* isolates samples sequenced in 2021. *flaA* (flagellin), a VF gene product identifying *C. jejuni*, was present in *S. enterica* isolates samples from human stool sampled in 2022 and all *C. jejuni* isolates samples, but not in *S. enterica* isolates samples from chicken meat, chicken stool, or human stool sampled in 2021.

Furthermore, certain VF genes such as *spvC* (type III secretion system effector *SpvC* phosphothreonine lyase) and pefB (plasmid-encoded fimbriae regulatory protein), associated with *S. enterica subsp. enterica* ser. Typhimurium str. LT2, were exclusively found in *S. enterica* isolates samples from human stool sampled in 2022. Conversely, *fyuA*, a pesticin/yersiniabactin receptor protein that identifies *Yersinia pestis*, was detected in every *S. enterica* isolates sample except those from human stool sampled in 2022. Finally, some VF genes, like *flif*, a flagellar M-ring protein known in *Yersinia enterocolitica* subsp. *enterocolitica*, were found in all samples, irrespective of the pathogen species.

#### Allele-based Pathogen Identification

The 19 *S. enterica* isolates samples were mapped against the reference genome of the expected pathogen, *S. enterica subsp. enterica* ser. Typhimurium (NC_003197.2 [74]), and the 65 *C. jejuni* isolates samples were mapped against *C. jejuni* (NC_002163.1).

The 19 *S. enterica* isolates samples have an average mapping coverage of 94.6% and an average mean mapping depth of 31 per base. The average total number of variants found per *S. enterica* isolates sample was 43,420. For the 65 *C. jejuni* isolates samples, the average mapping coverage was 93.7%, the average mean mapping depth was 42 per base and the average total number of variants found per sample was 26,654. These high values for the average total number of variants identified for samples were expected since the used subspecies for the mapping are different than the subspecies of the samples.

#### PathoGFAIR Samples Aggregation and Visualisation

The isolated samples exhibited a higher count of identified AMR genes compared to the metagenomic samples without prior isolation, enabling the incorporation of additional genes into concatenated gene consensus sequences. The resulting phylogenetic tree, constructed based on the AMR genes (Supplementary Figure S8), distinctly delineated different *S. enterica* subspecies. Similarly, this differentiation is evident in the phylogenetic tree based on the VF genes.

#### Sensitivity

Known species were successfully identified and confirmed across all workflows, including taxonomy profiling and VF genes identification (Supplementary Online Figure S6), achieving 100% expected pathogen detection at the expected taxonomic rank for all 84 tested samples resulting in a sensitivity of 1. Since there were no true negatives (TN) or conditions without pathogens in the dataset, specificity could not be calculated.

### Benchmarking PathoGFAIR

To evaluate the effectiveness of PathoGFAIR workflows, a benchmarking analysis was performed comparing PathoGFAIR’s pathogen detection capabilities with the systems and pipelines listed in Table 1. The primary goal was to assess each pipeline’s ability to accurately detect and identify pathogens from shotgun Nanopore metagenomic data using the samples without prior pathogen isolation. The detailed procedures for the selection and benchmarking process can be found on Protocols.io [78]. Replication instructions, including all benchmark systems, sample metadata, notebooks, and results, are publicly available through the PathoGFAIR GitHub repository.

### Selection of the pipelines for the benchmarking

To identify suitable systems and pipelines for the benchmark, each pipeline in Table 1 was evaluated based on availability, accepted input sequencing technique compatibility, and pathogen identification capability (Supplementary Table T4). Pipelines were classified as “Free Access” if available at no cost, “Free Trial” if partially or temporarily accessible, “Paid Access Only” if requiring payment, or “Non-functional” if outdated or inoperative. Compatibility with single-end Nanopore metagenomic sequencing data was verified to ensure each pipeline’s applicability for pathogen-focused workflows. Each pipeline’s pathogen identification capabilities were also assessed, examining relevant algorithms and tools to determine their accuracy and sensitivity within metagenomic datasets.

Following these criteria, pipelines requiring paid access (e.g. OneCodex) or being non-functional, as well as those incompatible with Nanopore data (e.g., SURPI, Sunbeam, Innuendo, and PAIPline) or lacking robust pathogen identification features, were excluded from further analysis.

Each selected pipeline was subsequently set up and tested to confirm ease of use and reproducibility. During testing, Victors was found to be non-functional, and SURPI presented difficulties, requiring local downloads of large host and pathogen reference databases. SURPI’s fragmented documentation contained outdated and contradictory requirements, with no updates since 2014. Innuendo, though documented, also presented usability challenges with incomplete setup instructions. BugSeq, while user-friendly and accessible through its web interface, had an average processing time of four hours from sample upload to results delivery. However, its free trial is limited to just 10 samples, and it lacked transparency regarding the tools, workflows, and databases used for analysis, as it is neither open-source nor adaptable. Ultimately, only IDseq (CZID) and our workflows, PathoGFAIR, met the selection criteria and were included for benchmarking (Supplementary Table T4).

IDseq offers a highly user-friendly interface, similar to PathoGFAIR on the Galaxy platform. However, unlike PathoGFAIR, IDseq’s pipeline is limited in terms of adaptability to its workflows, particularly with respect to the tools and parameters used. This restriction can hinder customisation and flexibility for users with specific analytical needs, reducing the pipeline’s versatility compared to PathoGFAIR. The IDseq pipeline took one hour for sample uploads and an additional hour and a half to complete the analysis across all samples.

### Benchmarking on samples without prior pathogen isolation

The benchmark was conducted using the 46 samples without prior pathogen isolation, from the first use case. The samples are chicken meat spiked with pathogens and sequenced using Nanopore as explained in [71]. Metadata for the 46 samples is summarised in Supplementary Table T1, including the expected *S. enterica* subspecies (*S. enterica* subsp. *houtenae* DSM 9221, *S. enterica* subsp. *enterica* DSM 554, and/or *S. enterica* subsp. *salamae* DSM 9220). For PathoGFAIR, taxonomy profiling from Workflow 2 and VF gene identification from Workflow 3 were evaluated.

In both PathoGFAIR and IDseq (Supplementary Table T3), *Salmonella enterica* was detected in all samples except one, resulting in a 97.8% detection rate. In the exceptional sample, *Salmonella* was not detected by either system. PathoGFAIR provided additional resolution, identifying the subspecies taxonomic rank in 60.9% of samples and the strain taxonomic rank in 13% of samples. In contrast, IDseq’s output report did not provide pathogen information beyond the species rank (Supplementary Figure S9).

Further benchmarking with diverse public metagenomic datasets could offer more comprehensive insights into the performance of PathoGFAIR across different experimental conditions and sample types. Such analyses would help assess the workflow’s robustness and adaptability to a wider range of use cases, contributing to its ongoing validation and refinement.

## Conclusion

In conclusion, we present PathoGFAIR, a collection of Galaxy FAIR adaptable workflows, designed for pathogen detection and tracking among samples. These five workflows span the entire analysis pipeline, ranging from preprocessing reads to advanced analyses including taxonomy profiling, virulence and antimicrobial resistance gene identification, SNP detection, and evolutionary history comparisons. The workflows generate diverse visualisations for a comprehensive understanding of the results, accompanied by interactive reports detailing all relevant inputs and outputs.

Our workflows have successfully identified pathogens down to genus, species, or subspecies taxonomic ranks across diverse samples, surpassing limitations observed in comparable pipelines. Our workflows facilitate comprehensive sample comparisons across diverse types, conditions, and sequencing techniques by offering interpretative and publication-ready visualisations. The open-access and user-friendlydesign of PathoGFAIR mitigates accessibilitychallenges and reduces reliance on local computational resources by leveraging Galaxy’s infrastructure for computational tasks, a feature that sets it apart from similar pipelines. This scalable workflow is a versatile solution for processing (meta)genomic samples, extending its utility beyond detecting foodborne pathogens.

In our findings, optimising sampling, preparation, and sequencing conditions, such as a 24-hour sample incubation, significantly enhances the identification of virulence and antimicrobial resistance genes. Indeed, the workflows’ performance correlates with sample characteristics, with higher CFU/mL values and read counts, and lower Ct values yielding more comprehensive results, which can be used to establish sampling guidelines. Moreover, as the Preprocessing workflow effectively removes host sequences, adaptive sampling during sequencing to exclude host DNA is not necessary. The workflows were still able to detect pathogens at least at species taxonomic rank for samples without prior pathogen isolation.

The experimental setup for this study serves as a proof of concept, demonstrating the feasibility of using WGS for the detection and characterisation of *S. enterica* in spiked food matrices. By detecting pathogens across varying CFU/mL levels, as shown in the spiking experiments, the workflows showcase their sensitivity and practical applicability in adhering to stringent regulatory standards. These results establish a solid foundation for applying PathoGFAIR in food safety laboratories and outbreak investigations, where detailed subspecies-rank information is critical for monitoring and traceability.

PathoGFAIR’s utility extends to enrichment-based analyses of foodborne pathogens, aligning with EU food safety standards such as EN/ISO 6579. Traditionally, these standards require enrichment followed by classical microbiological methods or PCR confirmation. PathoGFAIR complements these approaches by enabling direct analysis of enrichment broths through WGS, facilitating efficient detection and subspecies characterisation of *S. enterica*. This capability enhances food safety monitoring and outbreak response by differentiating *S. enterica* subspecies from previous outbreak strains, thereby advancing traceability and improving public health interventions.

We further supported the scientific community by introducing new 46 benchmark samples, making them publicly available. This demonstrates our significant investment of time and resources, providing valuable assets for future research.

In addition to the allele-based pathogen identification method, our workflow can be further enhanced by incorporating MLST. MLST, or Multi-Locus Sequence Typing, offers an alternative approach by characterising isolates through the sequences of housekeeping genes [49]. This method provides valuable information about the genetic diversity and evolutionary relationships among isolates, allowing for more precise identification and tracing of pathogens. By integrating MLST using MLST (v 2.22.0) tool [79]into our workflow, users can benefit from a comprehensive analysis that combines both alleles and variants identification methods, providing a more robust and accurate pathogen detection and tracing solution.

To address the complexity of detecting virulence and AMR genes located on mobile genetic elements (MGEs), future versions of PathoGFAIR can incorporate additional validation steps. Specifically, virulence and AMR gene detection could be cross-referenced with broader taxonomic markers (e.g., 16S rRNA) to ensure genes detected from MGEs are correctly attributed to their respective pathogenic hosts. These enhancements aim to further improve the accuracy of pathogen detection by more reliably linking virulence and AMR genes to their bacterial hosts, thereby refining the overall precision of the workflows.

In the future, integrating PathoGFAIR with Galaxy’s automated bot system holds the promise of ongoing updates and analyses requiring minimal human involvement. By establishing a dedicated bot for PathoGFAIR, continuous results will be effortlessly refreshed whenever new datasets are uploaded, similar to the Galaxy bot created for SARS-CoV-2 [80]. The Galaxy bot for SARS-CoV-2 automatically updates and reanalyses data with each new upload, maintaining up-to-date results and reducing the need for manual intervention. This automation ensures real-time, efficient data processing and analysis, enhancing the workflow’s accuracy and timeliness. Leveraging the user-friendly interface of the Galaxy platform ensures accessibility for users of all computational skill levels, streamlining the entire process from sample upload to result interpretation with ease. This study not only presents a robust computational solution but also lays the groundwork for semiautomated, efficient, and user-friendly pathogen detection and tracking workflows.

## Additional Files

Supplementary Figure S1. Violin plot with the percentage of quality-controlled host reads detected and removed in samples with respect to adaptive sampling during sequencing (Host excluded or not) Samples Without Prior Pathogen Isolation Supplementary online Figure S2. Krone Pie Chart for the taxonomy profiling Samples Without Prior Pathogen Isolation Supplementary Figure S3. Bar chart for the total number of VF genes (orange) and AMR genes (blue) found in samples with respect to incubation duration before DNA isolation Samples Without Prior Pathogen Isolation Supplementary Figure S4. Phylogenetic tree, using the nucleotide evolution model; General Time Reversible (GTR) model with a CAT approximation for rate heterogeneity across sites [54], for the identified AMR genes Samples Without Prior Pathogen Isolation Supplementary Figure S5. Upset plot illustrating the intersections of different metadata categories, including sampling year, pathogen species, and the original host of the samples, highlighting common and unique attributes among the datasets Samples With Prior Pathogen Isolation

Supplementary online Figure S6. Krona Pie Chart for the taxonomy profiling Samples With Prior Pathogen Isolation Supplementary Figure S7. Violin plot for the total number of VFs and AMR genes Samples With Prior Pathogen Isolation Supplementary Figure S8. Phylogenetic tree, using the nucleotide evolution model; General Time Reversible (GTR) model with a CAT approximation for rate heterogeneity across sites [54], for the identified AMR genes Samples With Prior Pathogen Isolation Supplementary Figure S9. Heatmap with PathoGFAIR benchmark using the first Use Case datasets and compare it with similar analysis systems presented in Table 1

Supplementary Table T1. Metadata for Samples Without Prior Pathogen Isolation

Supplementary Table T2. Metadata for Samples With Prior Pathogen Isolation

Supplementary Table T3. PathoGFAIR and IDSeq pathogen detection results for Samples Without Prior Isolation at various taxonomic ranks

Supplementary Table T4. Benchmarking PathoGFAIR systems and pipelines evaluation

## Availability of Source Code and Requirements

Lists the following:

- Project name: PathoGFAIR https://usegalaxy-eu.github.io/PathoGFAIR/
- Workflows on public Galaxy servers: https://training.galaxyproject.org/training-material/workflows/embed.html?query=pathogfair
- Workflows (v 0.1) on WorkflowHub: https://workflowhub.eu/search?utf8=%E2%9C%93&q=pathogfair
- Workflows (v 0.1) on Dockstore: https://dockstore.org/search?organization=iwc-workflows&entryType=workflows&search=engy
- Tutorial: https://training.galaxyproject.org/training-material/topics/microbiome/tutorials/pathogen-detection-from-nanopore-foodborne-data/tutorial.html
- Data analysis home page: https://github.com/usegalaxy-eu/PathoGFAIR
- Operating system(s): Platform independent
- Other requirements: Account on a Galaxy server
- License: MIT license

## Data Availability

The raw sequence reads of the 46 samples without prior isolation are available on Sequence Read Archive (SRA) under BioProjects [81]. The protocol for the preparation of these samples is available on Protocols.io [71]. The workflows presented in the Methods section are available on Intergalactic Workflow Commission (IWC) and two workflow registries (Dockstore and WorkflowHub). The training material to understand, learn, and try the workflows is available on the Galaxy Training Network (GTN) [68]. The Jupyter notebook for additional visualisations and generating the figures of this paper is available in a GitHub repository [82]. Benchmarking PathoGFAIR protocol is available on protocols.io [78].

## Declarations

## List of Abbreviations

AMR: Antimicrobial resistance
API: Application Programming Interface
CFU: Colony-forming unit
Ct: Cycle Threshold
EFSA: European Food Safety Authority
EU: European Union
FAIR: Findable Accessible Interoperable Resulable
GTN: Galaxy Training Network
IWC: Intergalactic Workflow Commission
MLST: Multilocus sequence typing
NGS: Next-Generation Sequencing
QC: Quality Control
RKI: Robert Koch Institute
SNP: Single-nucleotide polymorphism
SRA: Sequence Read Archive
VF: Virulence Factor
VFDB: Virulence Factor database
WHO: World Health Organization
WGS: Whole Genome Sequencing.

## Funding

This research was supported by the Digital Life Science Call for Academia-Industry Collaborations under EOSC-Life funding [83]. Additionally, financial support was provided by the German Federal Ministry of Education and Research BMBF grant 031 A538A de.NBI-RBC and the Ministry of Science, Research and the Arts Baden-Württemberg (MWK) within the framework of LI-BIS/de.NBI Freiburg. This work was supported by the Programme d’Investissements d’Avenir (PIA), grant Agence Nationale de la Recherche, number ANR-11-INBS-0013.

### Author’s Contributions

Engy Nasr (E.N.) led the formal analysis, investigation, software development, validation, visualisation, and writing of both the original draft and the review and editing phases. E.N. also designed the workflows, tested them, created training materials, documented and published datasets, workflows, codes, protocols, and wrote the manuscript. Anna Henger (A.H.) supported conceptualisation, formal analysis, funding acquisition, investigation, project administration, validation, visualisation, and writing. A.H. also prepared and sequenced all project datasets, ensuring alignment of analysis results with lab preparation conditions. Björn Grüning (B.G.) supported conceptualisation, formal analysis, funding acquisition, investigation, project administration, and software development.

B.G. integrated new databases into analytical tools and maintained Galaxy tools used in PathoGFAIR. Paul Zierep (P.Z.) contributed to formal analysis, investigation, software development, supervision, validation, visualisation, and writing. P.Z. supervised the project and edited parts of the workflows and training materials. Bérénice Batut (B.B.) led the conceptualisation, funding acquisition, methodology, project administration, software development, and supervision. B.B. applied for the EOSC Life Industry Call Grant, designed the project’s main goals and guidelines, and managed and edited the full project. All authors read and approved the final manuscript.

## Acknowledgements

This research is made possible by the invaluable support of the entire Freiburg Galaxy team, Bioinformatics, University of Freiburg. The authors extend their special thanks to Wolfgang Maier and Mina Ansari for their technical expertise and academic guidance. We also appreciate the contributions of all the researchers who provided input to the project, with particular gratitude to Tobias Schindler for his exceptional assistance and to Peter van Heusden for his insightful contributions.

## Supplementary Material

**Figure 5.**
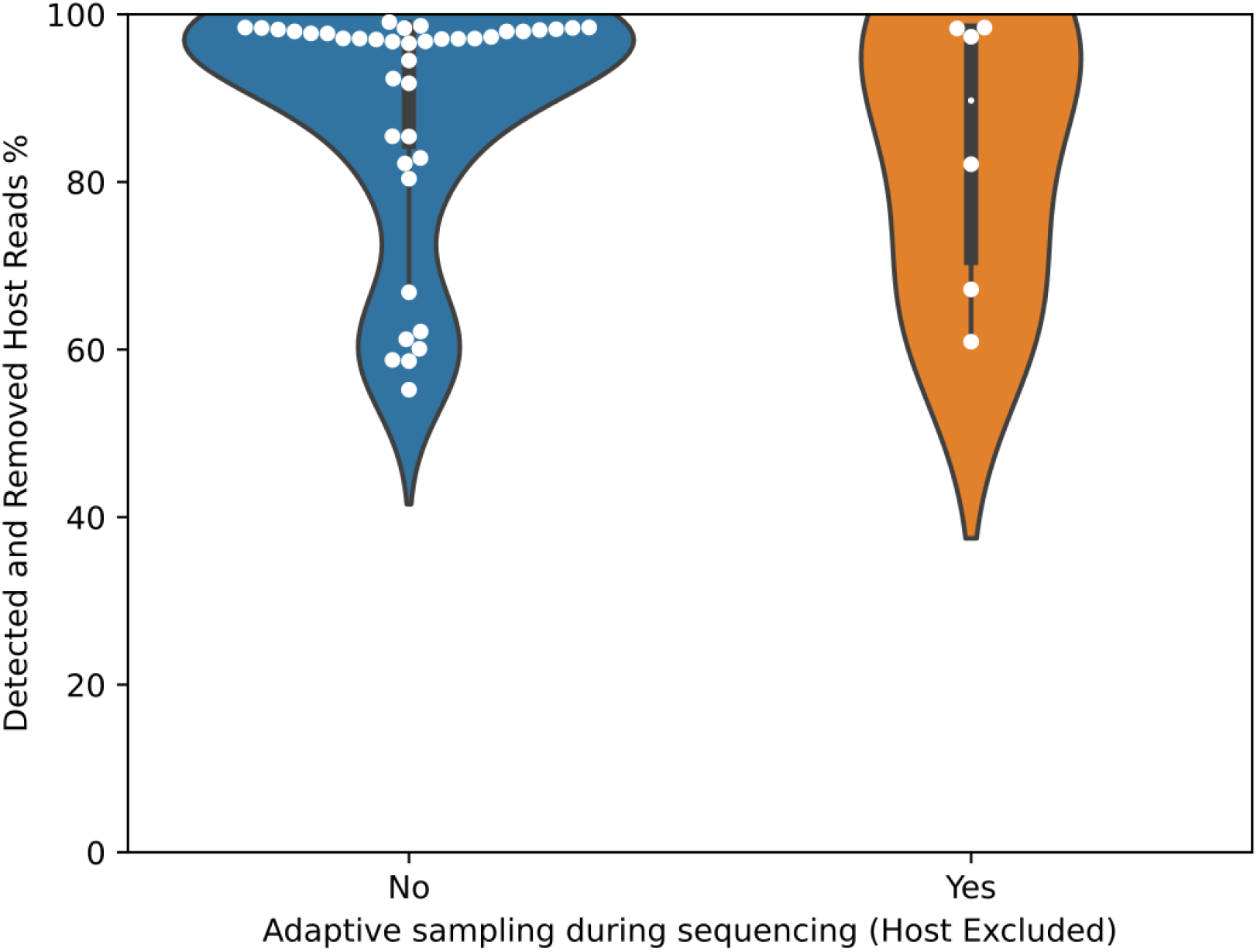
Supplementary Figure S1. Violin plot with the percentage of quality-controlled host reads detected and removed in samples with respect to adaptive sampling during sequencing (Host excluded or not) - Samples Without Prior Pathogen Isolation

**Figure 6.**
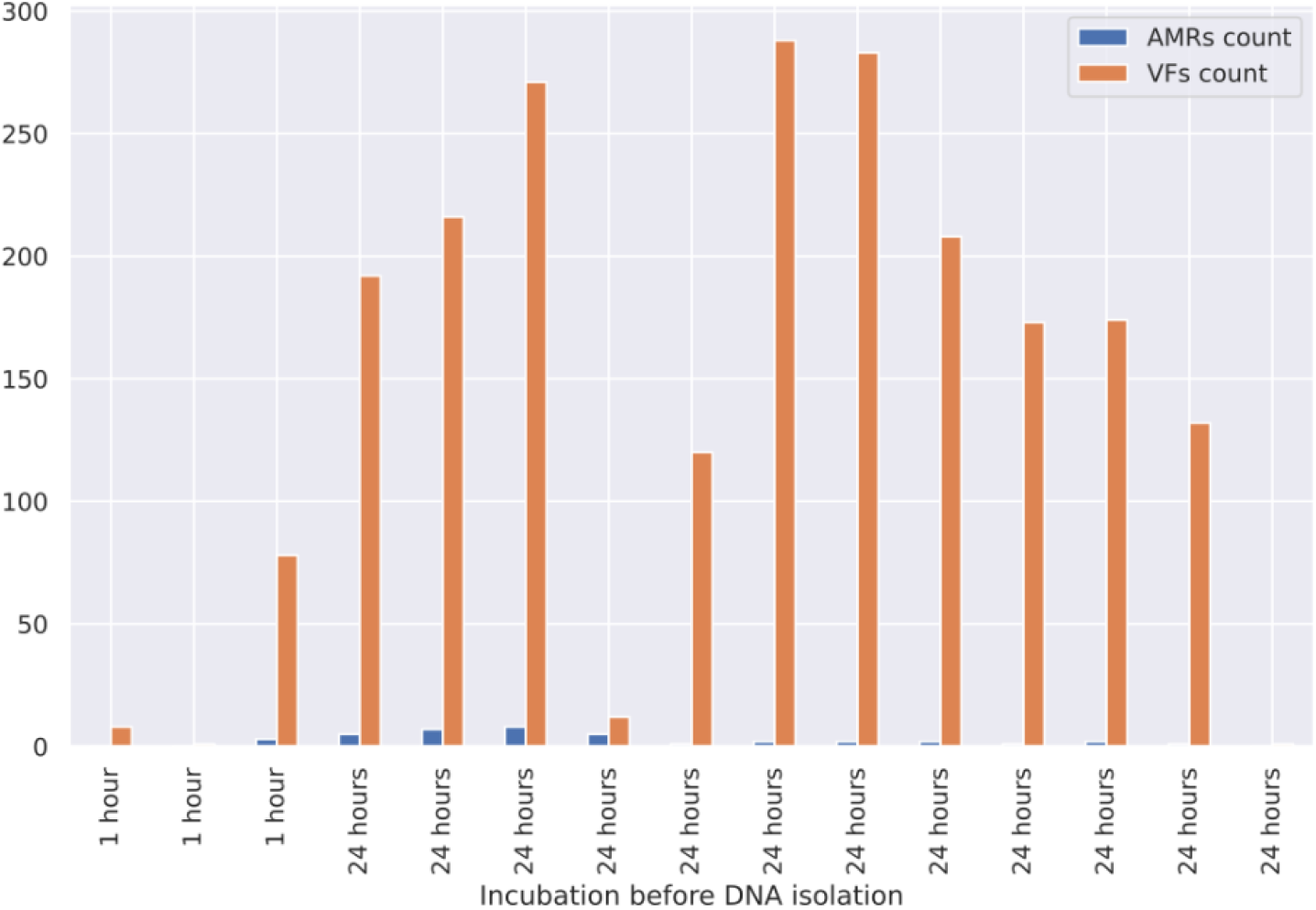
Supplementary Figure S3. Bar chart for the total number of VF genes (orange) and AMR genes (blue) found in samples with respect to incubation duration before DNA isolation - Samples Without Prior Pathogen Isolation

**Figure 7.**
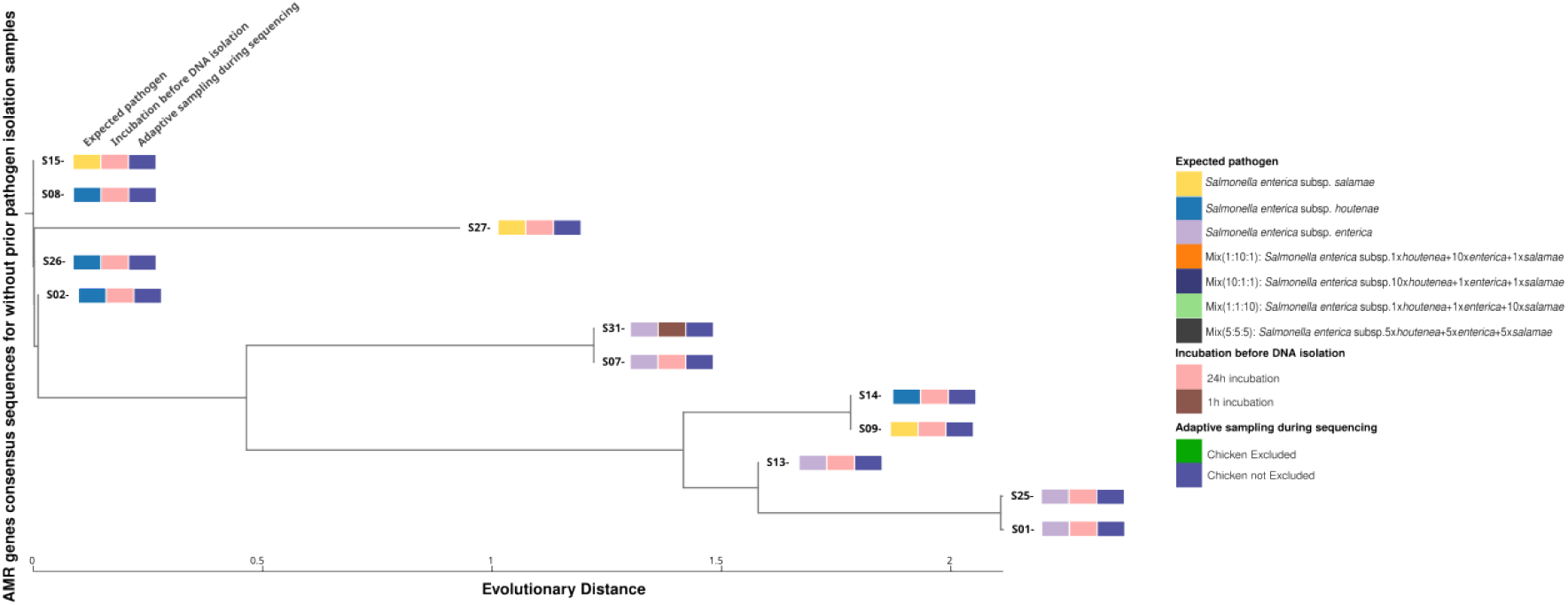
Supplementary Figure S4. Phylogenetic tree, using the nucleotide evolution model; General Time Reversible (GTR) model with a CAT approximation for rate heterogeneity across sites [54], for the identified AMR genes - Samples Without Prior Pathogen Isolation

**Figure 8.**
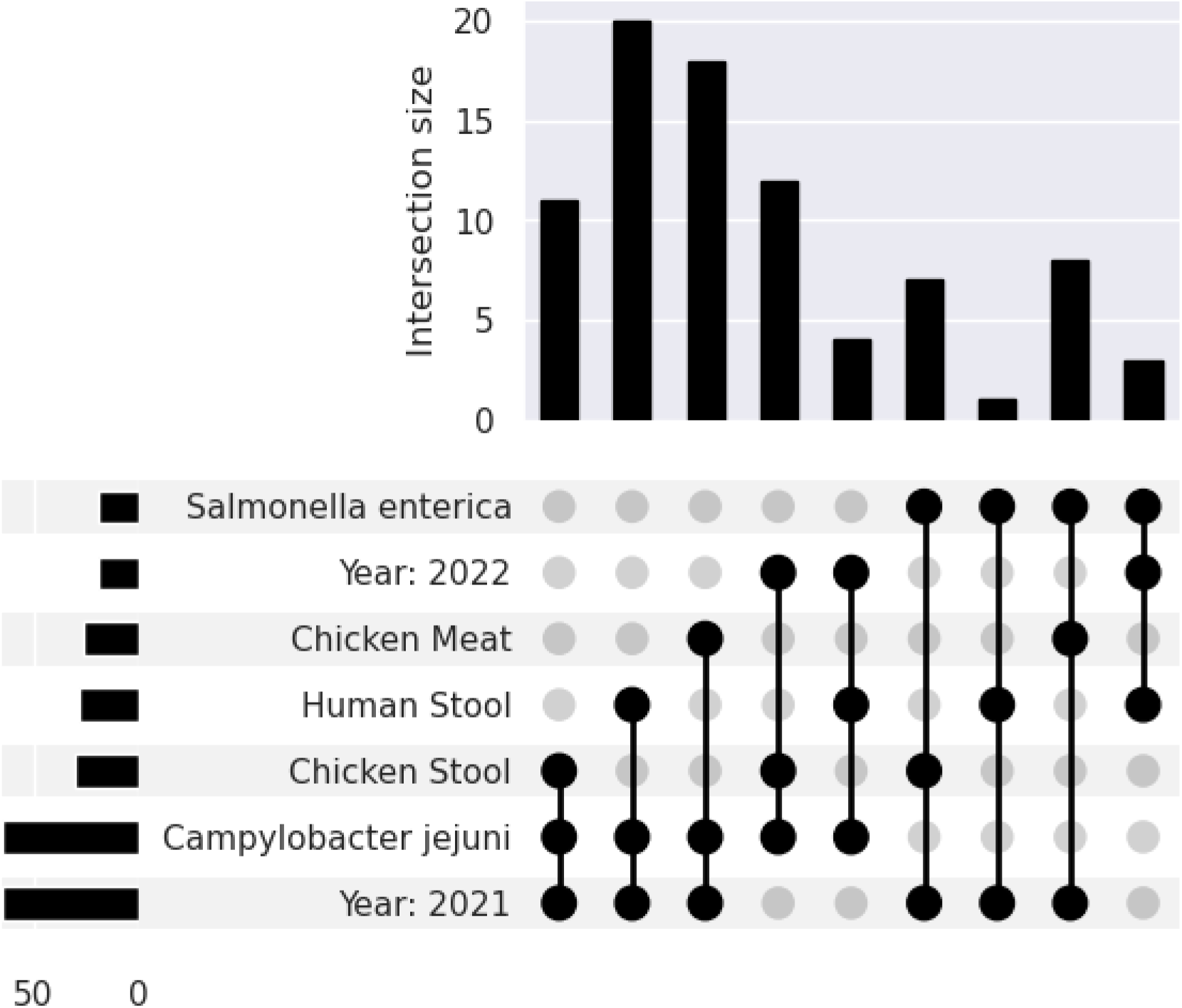
Supplementary Figure S5. Upset plot illustrating the intersections of different metadata categories, including sampling year, pathogen species, and the original host of the samples, highlighting common and unique attributes among the datasets - Samples With Prior Pathogen Isolation

**Figure 9.**
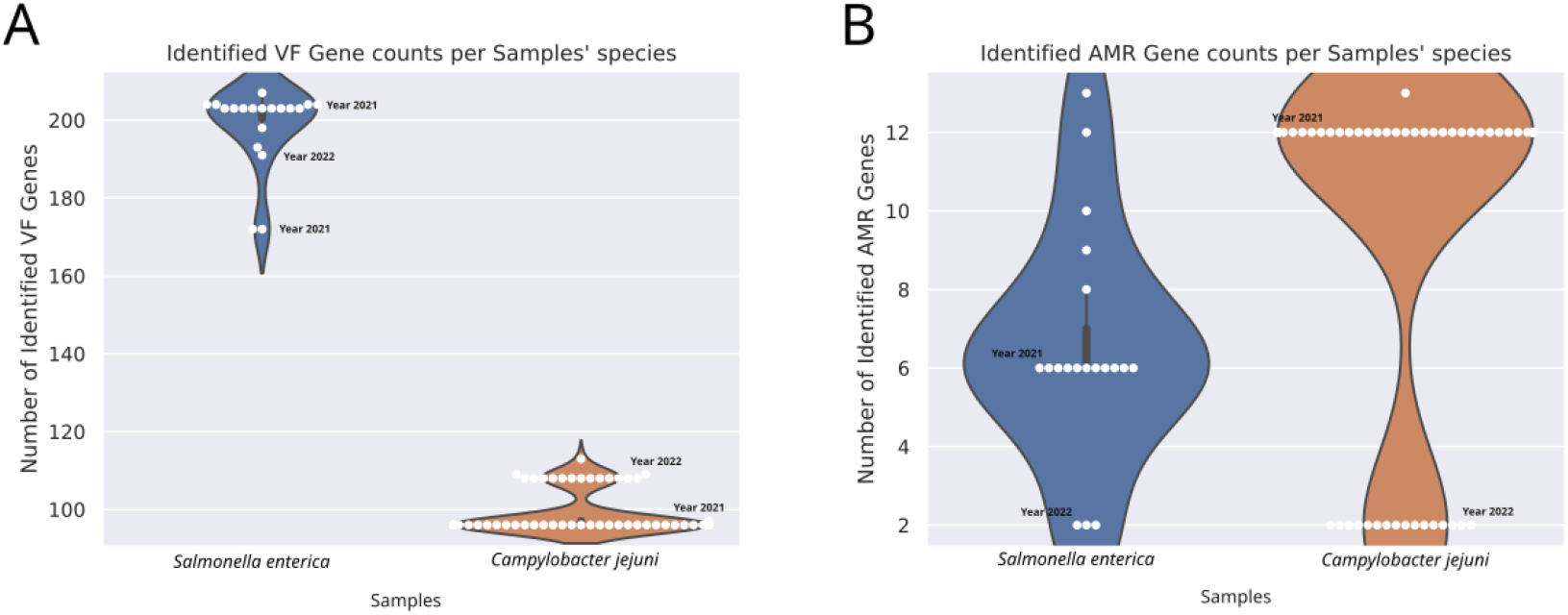
Supplementary Figure S7. Violin plot for the total number of VFs and AMR genes - Samples With Prior Pathogen Isolation

**Figure 10.**
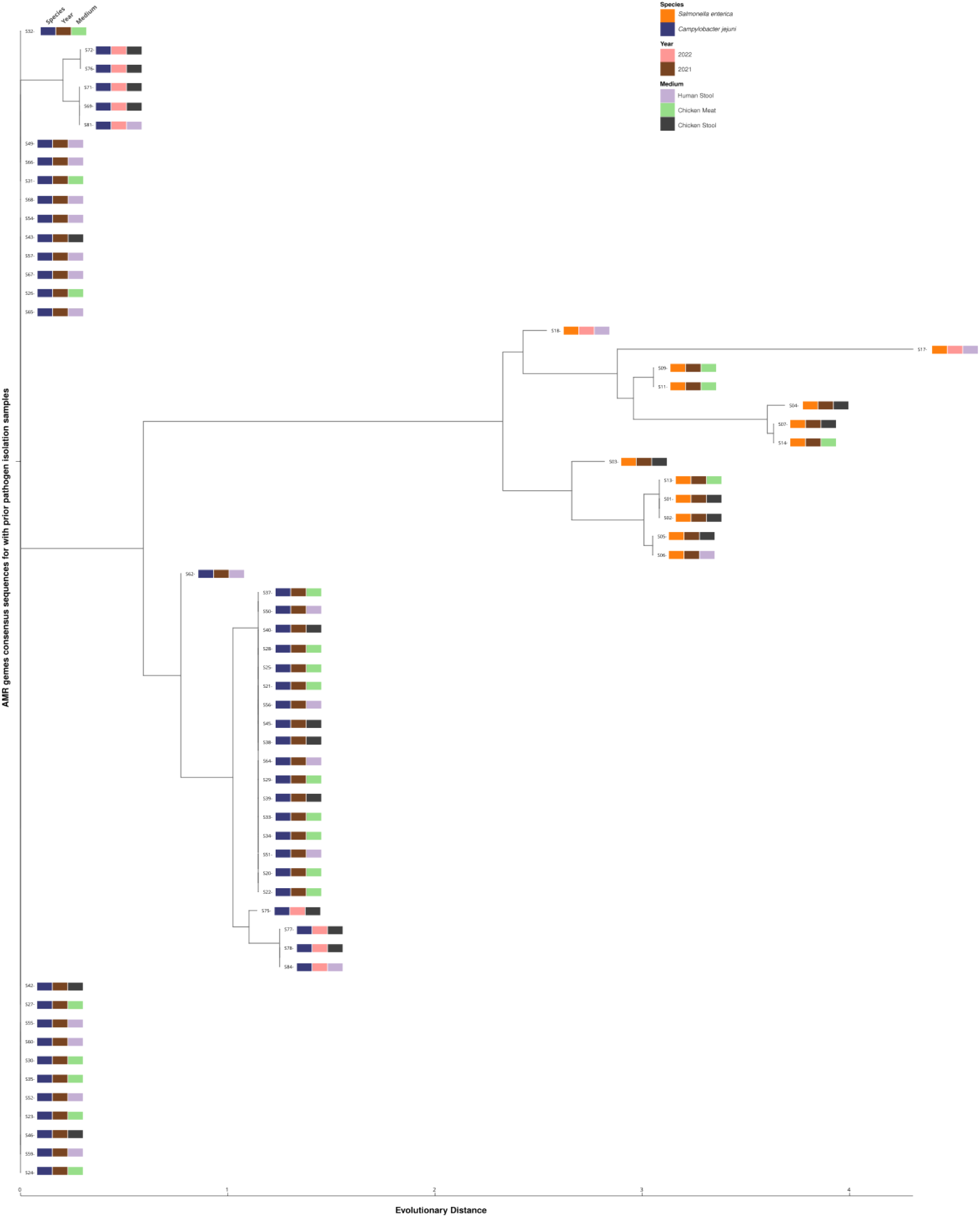
Supplementary Figure S8. Phylogenetic tree, using the nucleotide evolution model; General Time Reversible (GTR) model with a CAT approximation for rate heterogeneity across sites [54], for the identified AMR genes - Samples With Prior Pathogen Isolation

**Figure 11.**
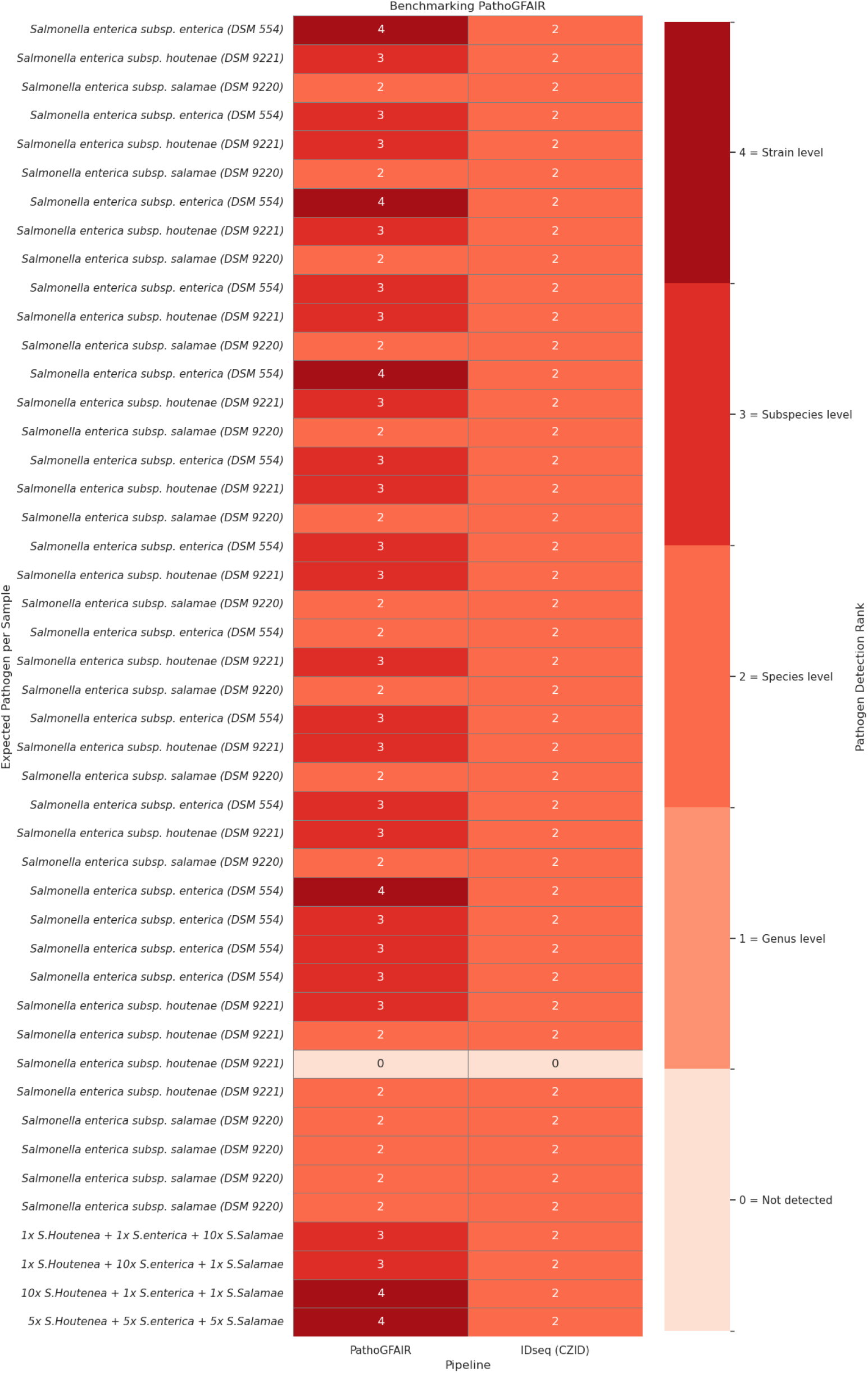
Supplementary Figure S9. Heatmap with PathoGFAIR benchmark using the first Use Case datasets and compare it with similar analysis systems presented in Table 1

